# Borderline *rpoB* mutations transmit at the same rate as common *rpoB* mutations in a tuberculosis cohort in Bangladesh

**DOI:** 10.1101/2023.02.14.528501

**Authors:** Pauline Lempens, Armand Van Deun, Kya J.M. Aung, Mohammad A. Hossain, Mahboobeh Behruznia, Tom Decroo, Leen Rigouts, Bouke C. de Jong, Conor J. Meehan

## Abstract

The spread of multidrug-resistant tuberculosis (MDR-TB) is a growing problem in many countries worldwide. Resistance to one of the primary first-line drugs, rifampicin, is caused by mutations in the *Mycobacterium tuberculosis rpoB* gene. While some of these infrequent mutations show lower fitness *in vitro* than more common mutations, their *in vivo* fitness is currently unknown.

We used a dataset of 394 whole genome sequenced MDR-TB isolates from Bangladesh, representing around 44% of notified MDR-TB cases over 6 years, to look at differences in transmission clustering between isolates with borderline *rpoB* mutations and those with common *rpoB* mutations. We found a relatively low percentage of transmission clustering in the dataset (34.8%) but no difference in clustering between different types of *rpoB* mutations. Compensatory mutations in *rpoA, rpoB*, and *rpoC* were associated with higher levels of transmission clustering as were lineages 2, 3, and 4 relative to lineage 1. Young people as well as patients with high sputum smear positive TB were more likely to be in a transmission cluster.

Our findings show that although borderline *rpoB* mutations have lower *in vitro* growth potential this does not translate into lower transmission potential or *in vivo* fitness. Proper detection of these mutations is crucial to ensure they do not go unnoticed and spread MDR-TB within communities.

**Data summary:** WGS reads are available in the European Nucleotide Archive (PRJEB39569). In addition, WGS reads, as well as pDST and clinical data, are included in the ReSeqTB data platform and are accessible on registration at https://platform.reseqtb.org/. Custom scripts for clustering are available at https://github.com/conmeehan/pathophy.

## Introduction

Tuberculosis (TB) is an airborne disease caused by *Mycobacterium tuberculosis* (MTB). It is one of the main causes of death from an infectious disease worldwide [1]. Antimicrobial drug resistance hinders control of the TB pandemic. On average, about 3-4% of newly diagnosed TB patients, and about 18-21% of patients previously treated for TB have rifampicin-resistant TB (RR-TB) or multidrug-resistant TB (MDR-TB, resistant to rifampicin and isoniazid) [1]. Although rifampicin resistance testing coverage and RR/MDR-TB treatment success rates have increased in the last decade, the global RR/MDR-TB burden remains unchanged [1]. Continued efforts to improve our understanding and control of RR/MDR-TB are therefore required.

RR/MDR-TB was long believed to occur mainly through acquisition of resistance during first-line treatment [2]. Molecular studies have however shown that the vast majority of RR/MDR-TB is transmitted [2]. In recent years, whole genome sequencing (WGS) has added a new dimension to (RR/MDR-)TB epidemiological research, offering the opportunity to investigate the dynamics and determinants of (RR/MDR-)TB transmission in greater detail, thereby providing insights on its optimal control [3, 4].

By definition, RR/MDR-TB isolates have deleterious mutations in the RNA polymerase (*rpoB*) gene, an essential gene for MTB. Not surprisingly, Ser450Leu, the most common *rpoB* mutation in MTB worldwide is also the one causing the least fitness loss, often accompanied by compensatory mutations in *rpoA* or *rpoC* [5, 6]. Less fit *rpoB* mutants tend to show *in vitro* growth defects that may lead to false rifampicin susceptibility in liquid based phenotypic drug susceptibility testing (pDST), while *in vivo* causing similarly high mortality and poor treatment outcome as Ser450Leu [7–10]. These so-called borderline mutations may spread more extensively in settings where the population is weakened, such as by HIV, further facilitated by inaccurate diagnostics not recognizing these strains as rifampicin resistant [11, 12]. We tested whether borderline mutations are as transmissible as ‘common’ *rpoB* mutations (i.e. similar transmission fitness) in a setting with low HIV co-infection where modern and ancestral lineages co-circulate.

## Materials and methods

### Study population

MDR-TB patients from the Damien Foundation MDR-TB project area in Bangladesh, which covers 13 of 64 districts of the country, diagnosed between 2005 and 2011, treated with a standardized short treatment regimen, and with WGS data of their baseline *M. tuberculosis* isolate available were included. Patients with an *rpoB* wild type baseline isolate or a baseline isolate containing only (an) *rpoB* mutation(s) with unknown association with rifampicin resistance (not listed in the “WHO Catalogue of mutations in *Mycobacterium tuberculosis* complex and their association with drug resistance” and located outside the *rpoB* rifampicin resistance-determining region (RRDR) (codons 426-452)), were excluded [13].

### Whole genome sequencing

Most isolates (367/394) were included in our earlier publications about the Bangladesh MDR-TB cohort, and their WGS was described there (35 in Lempens et al., 2018 and 332 in Lempens et al., 2020) [14, 15]. The remaining 27 isolates were sequenced according to the same procedure as described in Lempens et al., 2020, with WGS done at the Translational Genomics Research Institute through the ReSeqTB sequencing platform [14, 16]. Non-MTB reads as identified by Centrifuge were removed and isolates with >10% non-MTB reads were excluded [17]. Quality control of the MTB reads was done using the MTBseq pipeline and isolates with <90% coverage of the reference genome or an average sequencing depth <30x were excluded [18]. Command line version 2.8.12 of TBProfiler was used for read trimming and mapping, and variant calling and annotation [19, 20]. The literature-based TBProfiler library database (https://github.com/jodyphelan/tbdb) was accessed on 15 July 2020. In case of heteroresistance, variants of any frequency were included in the analysis. Drug resistance-associated variants with a sequencing depth below the default threshold of 10x but greater than 1x were included as well.

### Transmission clustering approach

Transmission clusters were calculated using a 5-SNP cut-off and a so-called loose (single linkage) clustering approach, in which isolates within a cluster had a maximum difference of 5 SNPs with at least one other isolate in the cluster [21]. To ensure only MDR-TB transmission was observed, clusters were further divided based on *rpoB* mutations, using an in-house developed Python script [22]. This meant that each transmission cluster contained only isolates with the same *rpoB* mutation, representing likely transmission of MDR-TB. If an isolate had one or more *rpoB* mutations in addition to the *rpoB* mutation shared with other isolates in the cluster, it was kept in the cluster. Custom scripts for clustering are available at https://github.com/conmeehan/pathophy.

### rpoB mutation classification

Based on their *rpoB* mutations as reported by TBProfiler, isolates were divided into three groups: common, low-confidence and borderline. The “common *rpoB* mutations” group consisted of isolates with one or more mutation(s) classified as “associated with resistance” in the “WHO Catalogue of mutations in *Mycobacterium tuberculosis* complex and their association with drug resistance” [13]. The “low-confidence *rpoB* mutations” group contained isolates with one or more mutation(s) classified in the WHO catalogue as “associated with resistance interim” because insufficient evidence on their association with rifampicin resistance exists. Their association with resistance is therefore based on the expert rule that any mutation within the *rpoB* RRDR (codons 426-452) should be assumed to confer resistance [13]. Using the same rule, we also included isolates with RRDR mutations not listed in the catalogue in the “low-confidence *rpoB* mutations” group. Isolates with a combination of “associated with resistance interim” or “RRDR” mutations and “associated with resistance” mutations (except 450Leu) were also included in this group. The third group, the “borderline *rpoB* mutations” group, consisted of isolates with a borderline mutation, alone or in combination with a second mutation. Based on the WHO catalogue, the following seven mutations were considered as borderline: Leu430Pro, Asp435Tyr, His445Asn, His445Leu, His445Ser, Leu452Pro, and Ile491Phe [13].

### Compensatory mutations

Mutations in *rpoA, rpoB*, and/or *rpoC* have been shown to compensate for the fitness loss associated with rifampicin resistance-related mutations in *rpoB* [23]. We looked for the link between such mutations and transmission clustering in our dataset. We created two definitions of potential compensatory mutations: any mutation in *rpoA* or *rpoC* and those in *rpoA/B/C* that have been confirmed as compensatory by Gygli et al., 2021 [24]. These mutations were determined from the Called files output from MTBseq.

### Statistics

Statistical analyses were carried out in R [25]. Multivariable logistic regression was carried out to investigate the association between type of *rpoB* mutation and presence in an MDR-TB transmission cluster. Other variables included were gender, age, sputum smear microscopy, lineage, fluoroquinolone resistance, and presence of a compensatory mutation. The model was simplified until all remaining variables were variables of interest or significantly associated with clustering. Significance p-value was set at 0.05.

### Ethics

All patients provided written informed consent prior to starting the shorter MDR-TB treatment regimen. The Institute of Tropical Medicine institutional review board provided ethics approval for the present deidentified analysis.

## Results

Between 2005 and 2011, 894 patients were notified as diagnosed with MDR-TB in the Damien Foundation MDR-TB project area. Of 894 patients, 581 started with a gatifloxacin-based standardized short treatment regimen, and 11/894 started with an ofloxacin-based regimen [26, 27]. For 414 patients, whole genome sequencing data of a baseline isolate was available. One patient was counted twice, as two treatment episodes were included, after confirmation of reinfection with a strain that was not in a transmission cluster with the strain of the first infection. Nineteen isolates were excluded because their *rpoB* gene was wild type and 1 isolate was excluded because the association between its *rpoB* mutation and rifampicin resistance was unknown and the mutation was located outside the RRDR. As a result, 394 isolates were included in the analysis (Supplementary Figure 1).

Based on the 5-SNP cut-off with loose clustering approach, 38.3% (151/394) of patients clustered into 43 transmission clusters, while 61.7% (243/394) of patients had a unique genotype. Of the 43 clusters, 6 clusters were divided based on *rpoB* mutations (Table 1). This resulted in 40 transmission clusters of isolates with the same *rpoB* mutation. Of 394 patients, 34.8% (137) were in one of these 40 transmission clusters, while 65.2% (257) of patients did not cluster. Clusters had a median size of 2.5 patients (range 2 to 16). In 2/40 clusters, one of the clustered isolates had a second *rpoB* mutation besides a Ser450Leu mutation shared within the cluster. In one case, this was a deletion within the RRDR (*rpoB*_c.1306_1308del) and in the other a Glu761Asp mutation (with unknown association with rifampicin resistance). In both cases, the cluster was not divided, but kept as such.

**Table 1.**
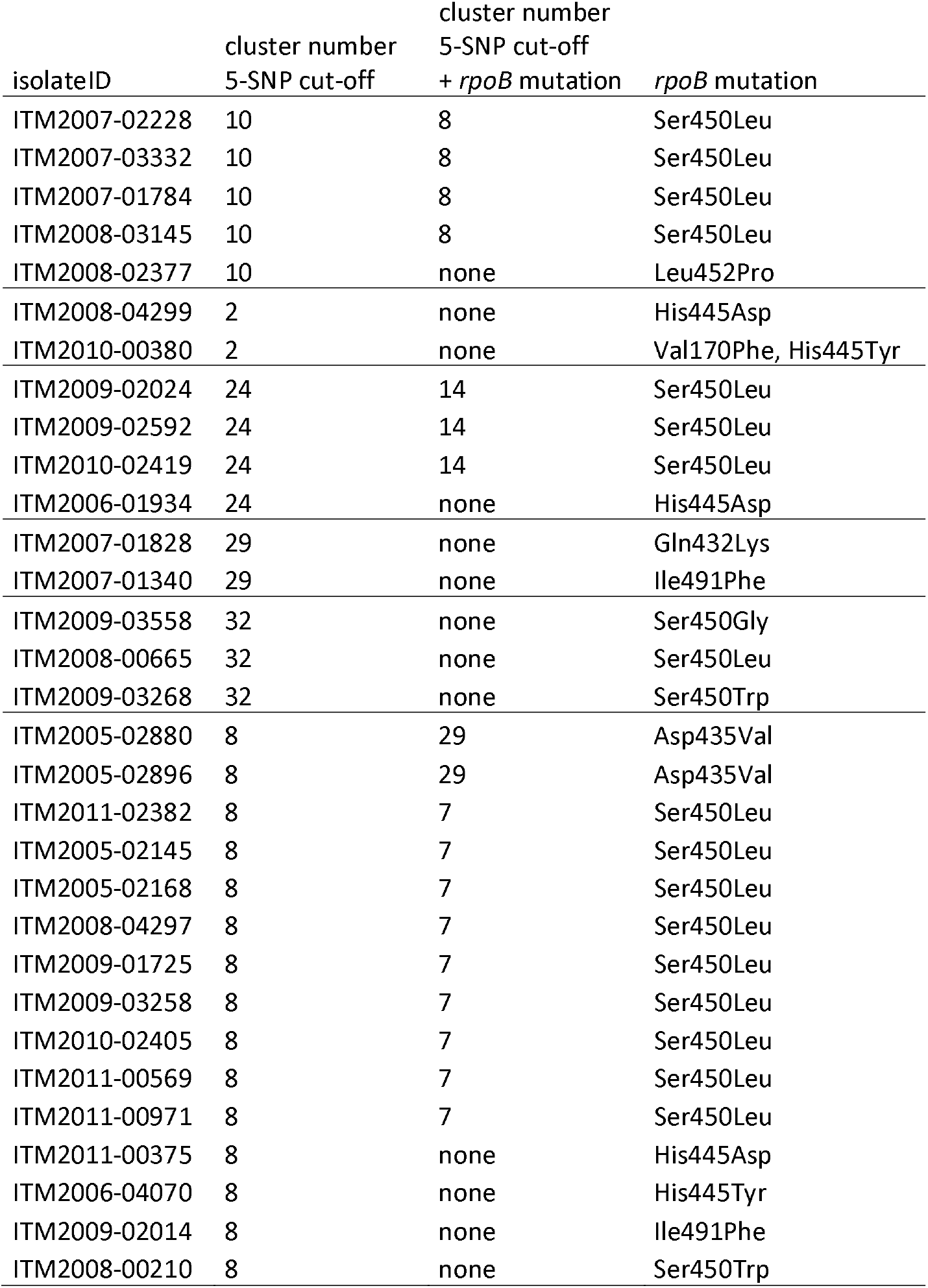
Overview of 5-SNP cut-off clusters that were split based on *rpoB* mutation.

| isolateID | cluster number<br>5-SNP cut-off | cluster number<br>5-SNP cut-off |  |
| --- | --- | --- | --- |
|  |  | + *rpoB* mutation | *rpoB* mutation |
| ITM2007-02228 | 10 | 8 | Ser450Leu |
| ITM2007-03332 | 10 | 8 | Ser450Leu |
| ITM2007-01784 | 10 | 8 | Ser450Leu |
| ITM2008-03145 | 10 | 8 | Ser450Leu |
| ITM2008-02377 | 10 | none | Leu452Pro |
| ITM2008-04299 | 2 | none | His445Asp |
| ITM2010-00380 | 2 | none | Val170Phe, His445Tyr |
| ITM2009-02024 | 24 | 14 | Ser450Leu |
| ITM2009-02592 | 24 | 14 | Ser450Leu |
| ITM2010-02419 | 24 | 14 | Ser450Leu |
| ITM2006-01934 | 24 | none | His445Asp |
| ITM2007-01828 | 29 | none | Gln432Lys |
| ITM2007-01340 | 29 | none | Ile491Phe |
| ITM2009-03558 | 32 | none | Ser450Gly |
| ITM2008-00665 | 32 | none | Ser450Leu |
| ITM2009-03268 | 32 | none | Ser450Trp |
| ITM2005-02880 | 8 | 29 | Asp435Val |
| ITM2005-02896 | 8 | 29 | Asp435Val |
| ITM2011-02382 | 8 | 7 | Ser450Leu |
| ITM2005-02145 | 8 | 7 | Ser450Leu |
| ITM2005-02168 | 8 | 7 | Ser450Leu |
| ITM2008-04297 | 8 | 7 | Ser450Leu |
| ITM2009-01725 | 8 | 7 | Ser450Leu |
| ITM2009-03258 | 8 | 7 | Ser450Leu |
| ITM2010-02405 | 8 | 7 | Ser450Leu |
| ITM2011-00569 | 8 | 7 | Ser450Leu |
| ITM2011-00971 | 8 | 7 | Ser450Leu |
| ITM2011-00375 | 8 | none | His445Asp |
| ITM2006-04070 | 8 | none | His445Tyr |
| ITM2009-02014 | 8 | none | Ile491Phe |
| ITM2008-00210 | 8 | none | Ser450Trp |

Of 394 isolates included, 322 had a common *rpoB* mutation, 13 had a low-confidence mutation, and 59 had a borderline mutation. (Combinations of) *rpoB* mutations found and their classification are presented in Supplementary Table 1.

The majority of patients were male (69.5%) and had a baseline isolate that was highly sputum smear positive (2+ or 3+) (88.6%) (Table 2). Isolates having a borderline *rpoB* mutation and isolates having a low-confidence mutation were found in a cluster at a similar frequency (35.6% and 38.5%, respectively) as isolates with a common *rpoB* mutation (34.5%). Of the 394 isolates, 51 were resistant to fluoroquinolones, of which 23 (45.1%) were found in a cluster. Four clusters contained one fluoroquinolone-resistant isolate each. One cluster contained two fluoroquinolone-resistant isolates, with different *gyrAB* mutations. Seven clusters contained at least two isolates with the same *gyrAB* mutation, suggesting transmission of pre-XDR TB.

**Table 2.** Factors associated with transmission clustering. Isolates within a transmission cluster had a maximum difference of 5 SNPs with at least one other isolate in the cluster and had the same *rpoB* mutation.

|  | Total<br>n | In a cluster<br>n (%) | Unique<br>n | Unadjusted OR<br>(95% CI) | Logistic regression |  |
| --- | --- | --- | --- | --- | --- | --- |
|  |  |  |  |  | Adjusted OR<br>(95% CI) | p value |
| Total | 394 | 137 (34.8) | 257 |  |  |  |
| Gender |  |  |  |  | NS |  |
| Male | 274 | 83 (30.3) | 191 | 1 |  |  |
| Female | 120 | 54 (45.0) | 66 | 1.9 (1.2-2.9) |  |  |
| Age |  |  |  |  |  |  |
| 10-24 | 124 | 71 (57.3) | 53 | 3.1 (2.0-5.1) | 2.8 (1.7-4.6) | <0.001 |
| 25-44 | 187 | 56 (29.9) | 131 | 1 | 1 |  |
| ≥45 | 83 | 10 (12.0) | 73 | 0.32 (0.15-0.64) | 0.33 (0.15-0.68) | 0.004 |
| Sputum smear |  |  |  |  |  |  |
| Negative/Scanty/1+ | 45 | 9 (20.0) | 36 | 1 | 1 |  |
| 2+/3+ | 349 | 128 (36.7) | 221 | 2.3 (1.1-5.3) | 2.3 (1.1-5.6) | 0.042 |
| Lineage |  |  |  |  |  |  |
| Lineage 1 | 115 | 17 (14.8) | 98 | 1 | 1 |  |
| Lineage 2 | 91 | 36 (39.6) | 55 | 3.8 (2.0-7.5) | 3.3 (1.6-6.9) | 0.001 |
| Lineage 3 | 69 | 24 (34.8) | 45 | 3.1 (1.5-6.4) | 2.5 (1.1-5.5) | 0.023 |
| Lineage 4 | 119 | 60 (50.4) | 59 | 5.9 (3.2-11.3) | 4.7 (2.4-9.6) | <0.001 |
| Fluoroquinolones |  |  |  |  | NS |  |
| Susceptible | 343 | 114 (33.2) | 229 | 1 |  |  |
| Resistant | 51 | 23 (45.1) | 28 | 1.7 (0.90-3.0) |  |  |
| Type of *rpoB* mutation |  |  |  |  |  |  |
| Common | 322 | 111 (34.5) | 211 | 1 | 1 |  |
| Low-confidence | 13 | 5 (38.5) | 8 | 1.2 (0.35-3.6) | 2.3 (0.62-8.3) | 0.190 |
| Borderline | 59 | 21 (35.6) | 38 | 1.1 (0.58-1.9) | 1.8 (0.90-3.5) | 0.096 |
| Any mutation in *rpoA/C* |  |  |  |  | NS |  |
| Absent | 290 | 98 (33.8) | 192 | 1 |  |  |
| Present | 104 | 39 (37.5) | 65 | 1.2 (0.73-1.9) |  |  |
| Confirmed<sup>#</sup><br>compensatory<br>*rpoA/B/C* mutation |  |  |  |  |  |  |
| Absent | 351 | 118 (33.6%) | 233 | 1 | 1 |  |
| Present | 43 | 19 (44.2%) | 24 | 1.6 (0.82-3.0) | 2.4 (1.1-5.0) | 0.025 |

<sup>#</sup> Confirmed by Gygli et al., 2021 [24]. NS: not significant

Isolates of modern *M. tuberculosis* lineage 2 (aOR 3.3; 95% CI 1.6-6.9), lineage 3 (aOR 2.5; 95% CI 1.1-5.5), and lineage 4 (aOR 4.7; 95% CI 2.4-9.6) clustered significantly more often than isolates of ancestral lineage 1 (Table 2). In addition, people aged 10-24 years clustered significantly more often than people aged 25-44 years (aOR 2.8; 95% CI 1.7-4.6), as did patients with highly sputum smear positive TB (2+ or 3+) compared to patients with a negative, scanty or 1+ sputum smear (aOR 2.3; 95% CI 1.1-5.6). People aged ≥45 were significantly less likely to be in a cluster relative to people aged 25-44 years (aOR 0.33; 95% CI0.15-0.68). Strains with a confirmed compensatory mutation (based on Gygli et al., 2021 [24]) were significantly more likely to be in a transmission cluster (aOR 2.4; 95% CI 1.1-5.0). Type of *rpoB* mutation, fluoroquinolone resistance, and presence of any mutations in *rpoA* or *rpoC* were not associated with being in a transmission cluster. Figure 1 shows a maximum likelihood tree of the isolates included in the study, with the clusters, as well as lineage and type of *rpoB* and compensatory mutation depicted in circles around the tree.

**Figure 1.**
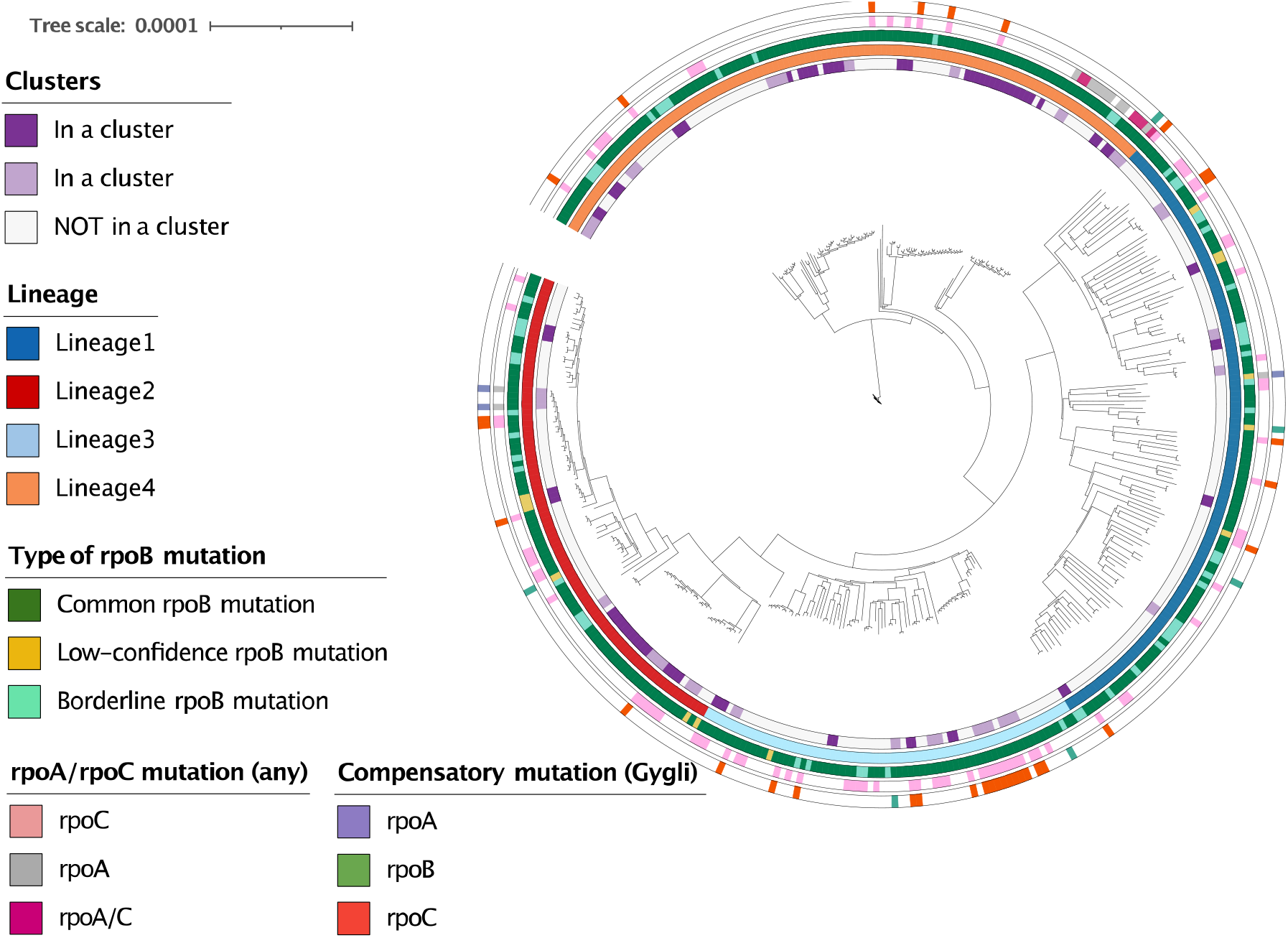
Maximum likelihood tree of the 394 isolates included in the study. Clusters, as well as lineage and type of *rpoB* and compensatory mutation are depicted in circles around the tree. Clusters are based on a 5-SNP cut-off combined with *rpoB* mutation, and are indicated in purple and lilac alternately to be able to distinguish adjacent clusters. Presence of any mutation in *rpoA* and/or *rpoC* is shown along with those in *rpoA/B/C* specifically found to be compensatory by Gygli et al., 2021 [24].

## Discussion

In our 2005-2011 MDR-TB cohort in the Bangladesh Damien Foundation MDR-TB project, 34.8% of MDR-TB resulted from transmission, with a median cluster size of 2.5 patients (range 2 to 16). Isolates with a borderline *rpoB* mutation and isolates with a low-confidence *rpoB* mutation clustered at a similar frequency (35.6% and 38.5%, respectively) as isolates having a common *rpoB* mutation (34.5%). Modern *M. tuberculosis* lineage (lineage 2, 3, or 4), young age, high sputum smear positivity, and compensatory mutations were associated with clustering.

Our results indicate that despite the *in vitro* fitness cost associated with borderline *rpoB* mutations, strains with such mutations seem to transmit equally well as strains with common *rpoB* mutations. In a drug-resistant TB cohort in Southern Brazil, 14/22 isolates (63.6%) with a borderline *rpoB* mutation were found in a cluster, compared to an overall clustering proportion of MDR-TB isolates of 73.4% [28].

We also find that compensatory mutations correlate positively with transmission clusters, irrespective of the underlying *rpoB* mutation. This suggests that these borderline mutations can transmit well, potentially with and without compensatory mutations, which should be accounted for in future studies. In contrast, labelling any mutation in *rpoA* or *rpoC* as compensatory did not correlate with transmission. This highlights that more understanding is needed into which specific mutations can compensate for fitness loss associated with antibiotic resistance as this is likely underestimated at present.

In addition to compensatory mutations, isolates with low-confidence or borderline *rpoB* mutations may benefit from obtaining a second *rpoB* mutation, with net advantage of increased rifampicin resistance compared to combined fitness loss [29]. We hypothesize that low-confidence or borderline *rpoB* mutations occur first if they occur sequentially in the same genome with a second *rpoB* mutation, rather than parallel in different bacilli. As a consequence, the effect on transmission could not have been present without the first sequential variant, characterised by the highest proportion mutant (unless both become fixed). Therefore, we included isolates with a combination of “associated with resistance interim” or “RRDR” mutations and “associated with resistance” mutations (except 450Leu) in the low-confidence group. Likewise, isolates with a combination of a borderline mutations and a second *rpoB* mutation were included in the borderline group.

Besides early and accurate detection of rifampicin resistance, effective treatment is crucial to stop RR/MDR-TB transmission. In the Bangladesh Damien Foundation MDR-TB project area, a standardized, highly effective treatment regimen has been in place for several decades now. In settings such as this, rifampicin resistance-associated variants that are more difficult to detect, such as the borderline *rpoB* mutations, may become the future TB endemic while variants that are detected and treated, are reduced.

The availability of effective RR/MDR-TB treatment in the Bangladesh Damien Foundation MDR-TB project area may explain the relatively low percentage of isolates that clustered. In a nationwide drug-resistant TB (DR-TB) surveillance study conducted in Bangladesh between 2011 and 2017, 38.4% of MDR-TB isolates were found in a cluster [30]. Our results are in line with these findings, despite the use of different transmission cluster estimation techniques (spoligotyping and 24-locus Mycobacterial Interspersed Repetitive Unit Variable Number Tandem Repeat vs WGS). Higher clustering percentages of MDR-TB isolates were reported in similar studies in Southern Brazil (73.4%), Tunisia (65.2%), and Saudi Arabia (67.6%), and of XDR-TB isolates in South Africa (85.5%) [22, 28, 31, 32]. In contrast, clustering percentages similar to the one found in our study were reported in Shenzhen (25.2%) and Shanghai (31.8%), in China [33, 34]. A caveat with transmission studies is that incomplete sampling fractions, as well as limited duration, tend to overestimate ‘unique’ isolates and thus underestimate TB due to recent transmission, as does migration into an area. While transmission to and from the area dense with garment manufacturers may not have been fully captured, the population based nature and 6 year study duration support that a low clustering rate may indeed be due to adequate detection and treatment of MDR-TB.

Lineages in the *Mycobacterium tuberculosis* complex are increasingly recognized to have co-evolved with humans, with recent lineages adapted to higher population densities where rapid spread is advantageous [35–37]. The modern lineages 2, 3, and 4 led to more secondary cases than the ancient lineages 1 and 6 [38, 39]. Our results suggest that known differences in transmissibility between *M. tuberculosis* lineages also apply to MDR-TB strains. In the DR-TB surveillance study in Bangladesh mentioned above, a significantly higher clustering rate was found in modern lineage isolates (49.3%) compared to ancient lineage isolates (9.6%) [30]. In an MDR-TB cohort in Saudi Arabia consisting of isolates belonging to lineage 1-4, no difference in clustering proportion was found between lineages [32]. However, since immigration rates in this setting were high (43.7% of patients was non-Saudi), transmission differences between lineages may not have been visible due to differences in lineage distribution between the highly diverse populations in the country. An alternative (partial) explanation for our findings could be the fact that lineage 1, and to a lesser extent lineage 2 have a higher molecular clock rate than lineage 4 [40, 41]. Using the same SNP cut-off for all lineages could therefore have led to an underestimation of transmission events of lineage 1 isolates compared to lineage 4 isolates [40].

The predominance of young people (age 10-24 years) in clusters is probably explained by recent industrialization in the Damien Foundation project area. Factories started to be built in large numbers around the country’s capital Dhaka not so long before the study period, and by now extend into the Southern part of the project area. From the project’s main MDR-TB detection districts in Greater Mymensingh, young people moved to these factory areas for work. Overcrowded living conditions and lack of employment protection when reporting ill, leading to self-medication with over-the-counter antibiotics, promote not only transmission but also creation of drug-resistant tuberculosis [42].

Our study has a few limitations. Firstly, not all RR-/MDR-TB patients in the Damien Foundation project area were diagnosed, started treatment, and had whole genome sequencing data available (Supplementary Figure 1). This may have led to an underestimation of the true clustering rate. In addition, besides the explanatory factors included in this study, other factors known to be associated with transmission of TB such as co-morbidities and host malnutrition were not assessed [43]. HIV-infection prevalence in the study setting is very low (<1%). Transmission of borderline *rpoB* mutants was likely underestimated since rifampicin resistance in these isolates more often remained undetected in the pre-GeneXpert study period and thus were not subjected to WGS. In addition, whole genome sequencing was done from cultured isolates, which may have led to culture bias affecting mutations with a relatively high fitness cost. Finally, transmission estimation of many lineages is not as well defined as for lineage 4 [21], meaning small over or under estimations can occur. Extensive transmission studies in lineages 1 and 3 will help refine these estimates in the future.

Transmission fitness is a complex phenotype, without straight correlation between replicative (e.g. *invitro*) fitness and transmission (e.g. *in vivo*) fitness [29]. Future studies that include data on the fitness cost of individual *rpoB* mutations and combinations of mutations could shed more light on the effect of fitness cost on transmission.

In conclusion, isolates with a borderline *rpoB* mutation and isolates with a low-confidence *rpoB* mutation transmitted equally well as isolates having a common *rpoB* mutation in our cohort of MDR-TB patients from Bangladesh. In addition, modern *M. tuberculosis* lineage 2, 3, and 4 were associated with transmission. Our findings stress the importance of accurate detection of resistant variants in order to maximally limit their spread.

## Acknowledgements

Damien Foundation Belgium and the staff of the Damien Foundation Bangladesh TB/Leprosy control project have supported excellent and innovative DR-TB care for decades, including diagnostic analyses at the Institute of Tropical Medicine (ITM) Antwerp. Their meticulous programmatic documentation of extensive clinical information allowed for the present analysis. Our gratitude also goes to the laboratory technicians at the Unit of Mycobacteriology, ITM Antwerp for their excellent support. The sequencing work was performed under the direction of David Engelthaler at the Translational Genomics Research Institute through the ReSeqTB sequencing platform led by Marco Schito and supported by the Bill & Melinda Gates Foundation (OPP1115887).

## Author contributions

Conceptualization: PL, LR, BCdJ, CJM; Data curation: PL, AVD, KJMA, MAH; Formal Analysis: PL, MB, CJM; Funding acquisition: AVD, BCdJ, CJM; Investigation: PL, AVD, TD, CJM; Methodology: PL, AVD, TD, LR, BCdJ, CJM; Project administration: PL, AVD, KJMA, MAH; Resources: AVD, KJMA, MAH; Supervision: TD, LR, BCdJ, CJM; Visualization: PL, MB, CJM; Writing – original draft: PL, CJM; Writing – review & editing: PL, AVD, KJMA, MAH, MB, TD, LR, BCdJ, CJM

## References

1. WHO. Global tuberculosis report 2021. 2021; Available from: https://www.who.int/publications/i/item/9789240037021.

2. Dheda, K., et al., The epidemiology, pathogenesis, transmission, diagnosis, and management of multidrug-resistant, extensively drug-resistant, and incurable tuberculosis. Lancet Respir Med, 2017.

3. Comas, I., Genomic Epidemiology of Tuberculosis. Adv Exp Med Biol, 2017. 1019: p. 79–93.

4. Gagneux, S., Ecology and evolution of Mycobacterium tuberculosis. Nat Rev Microbiol, 2018. 16(4): p. 202–213.

5. Brandis, G. and D. Hughes, Genetic characterization of compensatory evolution in strains carrying rpoB Ser531Leu, the rifampicin resistance mutation most frequently found in clinical isolates. J Antimicrob Chemother, 2013. 68(11): p. 2493–7.

6. Comas, I., et al., Whole-genome sequencing of rifampicin-resistant Mycobacterium tuberculosis strains identifies compensatory mutations in RNA polymerase genes. Nat Genet, 2011. 44(1): p. 106–10.

7. Van Deun, A., et al., Rifampin drug resistance tests for tuberculosis: challenging the gold standard. J Clin Microbiol, 2013. 51(8): p. 2633–40.

8. Van Deun, A., et al., Disputed rpoB mutations can frequently cause important rifampicin resistance among new tuberculosis patients. Int J Tuberc Lung Dis, 2015. 19(2): p. 185–90.

9. Miotto, P., et al., Role of Disputed Mutations in the rpoB Gene in Interpretation of Automated Liquid MGIT Culture Results for Rifampin Susceptibility Testing of Mycobacterium tuberculosis. J Clin Microbiol, 2018. 56(5).

10. Torrea, G., et al., Variable ability of rapid tests to detect Mycobacterium tuberculosis rpoB mutations conferring phenotypically occult rifampicin resistance. Sci Rep, 2019. 9(1): p. 11826.

11. Makhado, N.A., et al., Outbreak of multidrug-resistant tuberculosis in South Africa undetected by WHO-endorsed commercial tests: an observational study. Lancet Infect Dis, 2018. 18(12): p. 1350–1359.

12. Loiseau, C., et al., HIV Coinfection Is Associated with Low-Fitness rpoB Variants in Rifampicin-Resistant Mycobacterium tuberculosis. Antimicrob Agents Chemother, 2020. 64(10).

13. WHO. Catalogue of mutations in Mycobacterium tuberculosis complex and their association with drug resistance. 2021; Available from: https://www.who.int/publications/i/item/9789240028173.

14. Lempens, P., et al., Initial resistance to companion drugs should not be considered an exclusion criterion for the shorter multidrug-resistant tuberculosis treatment regimen. Int J Infect Dis, 2020. 100: p. 357–365.

15. Lempens, P., et al., Isoniazid resistance levels of Mycobacterium tuberculosis can largely be predicted by high-confidence resistance-conferring mutations. Sci Rep, 2018. 8(1): p. 3246.

16. Starks, A.M., et al., Collaborative Effort for a Centralized Worldwide Tuberculosis Relational Sequencing Data Platform. Clin Infect Dis, 2015. 61 Suppl 3(Suppl 3): p. S141–6.

17. Kim, D., et al., Centrifuge: rapid and sensitive classification of metagenomic sequences. Genome Res, 2016. 26(12): p. 1721–1729.

18. Kohl, T.A., et al., MTBseq: a comprehensive pipeline for whole genome sequence analysis of Mycobacterium tuberculosis complex isolates. PeerJ, 2018. 6: p. e5895.

19. Coll, F., et al., Rapid determination of anti-tuberculosis drug resistance from whole-genome sequences. Genome Med, 2015. 7(1): p. 51.

20. Phelan, J.E., et al., Integrating informatics tools and portable sequencing technology for rapid detection of resistance to anti-tuberculous drugs. Genome Med, 2019. 11(1): p. 41.

21. Meehan, C.J., et al., The relationship between transmission time and clustering methods in Mycobacterium tuberculosis epidemiology. EBioMedicine, 2018. 37: p. 410–416.

22. Oostvogels, S., et al., Transmission, distribution and drug resistance-conferring mutations of extensively drug-resistant tuberculosis in the Western Cape Province, South Africa. Microb Genom, 2022. 8(4).

23. Gagneux, S., Fitness cost of drug resistance in Mycobacterium tuberculosis. Clin Microbiol Infect, 2009. 15 Suppl 1: p. 66–8.

24. Gygli, S.M., et al., Prisons as ecological drivers of fitness-compensated multidrug-resistant Mycobacterium tuberculosis. Nat Med, 2021. 27(7): p. 1171–1177.

25. R Core Team, R: A language and environment for statistical computing. 2020, R Foundation for Statistical Computing: Vienna, Austria.

26. Van Deun, A., et al., Short, highly effective, and inexpensive standardized treatment of multidrug-resistant tuberculosis. Am J Respir Crit Care Med, 2010. 182(5): p. 684–92.

27. Aung, K.J., et al., Successful ‘9-month Bangladesh regimen’ for multidrug-resistant tuberculosis among over 500 consecutive patients. Int J Tuberc Lung Dis, 2014. 18(10): p. 1180–7.

28. Salvato, R.S., et al., Genomic-based surveillance reveals high ongoing transmission of multi-drug-resistant Mycobacterium tuberculosis in Southern Brazil. Int J Antimicrob Agents, 2021: p. 106401.

29. Borrell, S. and S. Gagneux, Infectiousness, reproductive fitness and evolution of drug-resistant Mycobacterium tuberculosis. Int J Tuberc Lung Dis, 2009. 13(12): p. 1456–66.

30. Rahman, S.M.M., et al., Molecular Epidemiology and Genetic Diversity of Multidrug-Resistant Mycobacterium tuberculosis Isolates in Bangladesh. Microbiol Spectr, 2022. 10(1): p. e0184821.

31. Bouzouita, I., et al., Whole-Genome Sequencing of Drug-Resistant Mycobacterium tuberculosis Strains, Tunisia, 2012-2016. Emerg Infect Dis, 2019. 25(3): p. 538–546.

32. Al-Ghafli, H., et al., Drug-resistance profiling and transmission dynamics of multidrug-resistant Mycobacterium tuberculosis in Saudi Arabia revealed by whole genome sequencing. Infect Drug Resist, 2018. 11: p. 2219–2229.

33. Jiang, Q., et al., Citywide Transmission of Multidrug-resistant Tuberculosis Under China’s Rapid Urbanization: A Retrospective Population-based Genomic Spatial Epidemiological Study. Clin Infect Dis, 2020. 71(1): p. 142–151.

34. Yang, C., et al., Transmission of multidrug-resistant Mycobacterium tuberculosis in Shanghai, China: a retrospective observational study using whole-genome sequencing and epidemiological investigation. Lancet Infect Dis, 2017. 17(3): p. 275–284.

35. Gagneux, S., Host-pathogen coevolution in human tuberculosis. Philos Trans R Soc Lond B Biol Sci, 2012. 367(1590): p. 850–9.

36. Correa-Macedo, W., G. Cambri, and E. Schurr, The Interplay of Human and Mycobacterium Tuberculosis Genomic Variability. Front Genet, 2019. 10: p. 865.

37. Cardona, P.J., M. Català, and C. Prats, The Origin and Maintenance of Tuberculosis Is Explained by the Induction of Smear-Negative Disease in the Paleolithic. Pathogens, 2022. 11(3).

38. Coscolla, M. and S. Gagneux, Consequences of genomic diversity in Mycobacterium tuberculosis. Semin Immunol, 2014. 26(6): p. 431–44.

39. Walker, T.M., et al., Mycobacterium tuberculosis transmission in Birmingham, UK, 2009-19: An observational study. Lancet Reg Health Eur, 2022. 17: p. 100361.

40. Menardo, F., et al., The molecular clock of Mycobacterium tuberculosis. PLOS Pathogens, 2019. 15(9): p. e1008067.

41. Ford, C.B., et al., Mycobacterium tuberculosis mutation rate estimates from different lineages predict substantial differences in the emergence of drug-resistant tuberculosis. Nat Genet, 2013. 45(7): p. 784–90.

42. Bank, W. Bangladesh Social Protection Public Expenditure Review (PER). 2021; Available from: https://documents1.worldbank.org/curated/en/829251631088806963/pdf/Bangladesh-Social-Protection-Public-Expenditure-Review.pdf.

43. Lönnroth, K., et al., Drivers of tuberculosis epidemics: the role of risk factors and social determinants. Soc Sci Med, 2009. 68(12): p. 2240–6.

