## Supplemental Table 1 for "Borderline *rpoB* mutations transmit at the same rate as common *rpoB* mutations in a tuberculosis cohort in Bangladesh"

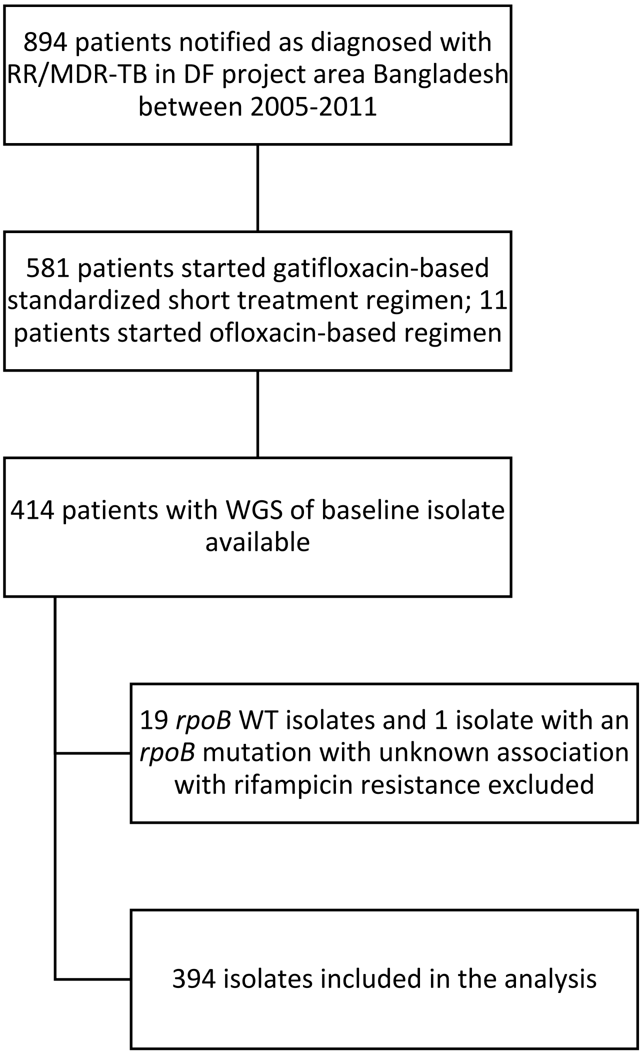


Supplementary Figure 1. Flowchart of sample selection.

Supplementary Table 1. (Combinations of) *rpoB* mutations found and their classification as "associated with resistance", "associated with resistance interim", or "borderline" in the "WHO Catalogue of mutations in *Mycobacterium tuberculosis* complex and their association with drug resistance", or as "not listed in the WHO catalogue but located within the RRDR", or "not listed in the WHO catalogue and not located within the RRDR (unknown association)" [13]. *rpoB* mutations were further grouped as "common", "low-confidence" or "borderline as described in the materials and methods section. Isolates within a transmission cluster had a maximum difference of 5 SNPs with at least one other isolate in the cluster and had the same *rpoB* mutation.

| Type of *rpoB* mutation | Total (n) | In a cluster (n) | Unique (n) |
|---|---|---|---|
| **Common** | | | |
| 1296_1297insTTC | 2 | 0 | 2 |
| 1296_1297insTTC, 1299_1300insTTG | 1 | 0 | 1 |
| Val170Phe | 5 | 0 | 5 |
| Val170Phe, His445Tyr | 1 | 0 | 1 |
| Gln432Lys | 3 | 0 | 3 |
| Gln432Pro | 2 | 0 | 2 |
| Asp435Val | 16 | 4 | 12 |
| Ser441Leu | 1 | 0 | 1 |
| His445Arg | 4 | 0 | 4 |
| His445Asp | 24 | 2 | 22 |
| His445Cys | 2 | 0 | 2 |
| His445Tyr | 30 | 7 | 23 |
| Ser450Leu | 210 | 92 | 118 |
| Ser450Leu, 1306_1308del | 1 | 1 | 0 |
| Ser450Leu, Ala286Val | 1 | 0 | 1 |
| Ser450Leu, Ile480Val | 1 | 0 | 1 |
| Ser450Leu, Glu761Asp | 1 | 1 | 0 |
| Ser450Trp | 17 | 4 | 13 |
| **Low-confidence** | | | |
| Thr427Pro, His445Tyr | 3 | 3 | 0 |
| Leu430Arg, Ser450Val | 1 | 0 | 1 |
| Ser431Arg, Asp435Gly, Asn437Asp | 1 | 0 | 1 |
| Gln432Arg | 1 | 0 | 1 |
| Met434Thr, Asp435Gly | 2 | 2 | 0 |
| Asp435Gly, His445Asp | 1 | 0 | 1 |
| His445Gln, Ser450Trp | 1 | 0 | 1 |
| His445Gly | 1 | 0 | 1 |
| His445Pro | 1 | 0 | 1 |
| Ser450Gly | 1 | 0 | 1 |
| **Borderline** | | | |
| Leu430Pro | 8 | 4 | 4 |
| Leu430Pro, Ser493Leu | 1 | 0 | 1 |
| Asp435Tyr | 17 | 7 | 10 |
| Asp435Tyr, Ser431Asn | 1 | 1 | 0 |
| Asp435Tyr, Met434Arg | 1 | 1 | 0 |
| His445Asn | 6 | 5 | 1 |
| His445Asn, Asp435Glu | 1 | 0 | 1 |
| His445Leu | 4 | 0 | 4 |
| His445Leu, Ser428Thr | 3 | 3 | 0 |
| Leu452Pro | 11 | 0 | 11 |
| Ile491Phe | 6 | 0 | 6 |
